# A robust intensity distribution alignment for harmonization of T1w intensity values

**DOI:** 10.1101/2022.06.15.496227

**Authors:** Donatas Sederevičius, Atle Bjørnerud, Kristine B. Walhovd, Koen Van Leemput, Bruce Fischl, Anders M. Fjell

## Abstract

Variations in image intensities between magnetic resonance imaging (MRI) acquisitions affect the subsequent image processing and its derived outcomes. Therefore, it is necessary to normalize images of different scanners/acquisitions, especially for longitudinal studies where a change of scanner or pulse sequence often happens. Here, we propose a robust intensity distribution alignment (RIDA) method to remove between-scan effects. The method is based on MRI T1w images acquired in close succession and robustly aligns two cumulative distribution functions (CDF) of voxel intensities to improve image-derived outcomes of a range of subcortical brain structures with different acquisition parameters. We compare RIDA with the other image harmonization methods: *mica* and RAVEL. We study three intra-scanner and three inter-scanner protocol variations among the same 20 participants scanned with Siemens 1.5T Avanto, 3T Skyra, and 3T Prisma scanners on the same day and use image-derived volumetric outputs from the Sequence Adaptive Multimodal Segmentation (SAMSEG) method. We find that CDF-based intensity harmonization (*mica* and RIDA) significantly reduces intensity differences, improves consistency in volume quantification, and increases spatial overlap between two images acquired in close succession. The improvements are most considerable if the intensity normalization is based on subcortical structures only (RIDA), excluding cortical regions, instead of the whole brain. However, the effect of the corrections varies considerably as a function of the compared scanners and sequences. In conclusion, the RIDA scan-effect normalization improves the consistency of image-derived measures, but its performance depends on several factors.

## 1. Introduction

Magnetic resonance imaging (MRI) parameters such as pulse sequence, scanner type, or RF coil configurations can alter signal-to-noise and contrast-to-noise ratios between images (Han et al., 2006; Jovicich et al., 2009). In addition, MRI images are collected in arbitrary units of intensity which are often not comparable across visits, even within the same subject and scanner. For instance, in the Alzheimer’s Disease Neuroimaging Initiative (ADNI) and Australian Imaging, Biomarkers and Lifestyle (AIBL) studies, which have highly standardized protocols, noticeable differences in the raw intensities are observed between imaging sites (Shinohara et al., 2014). Such intensity variations between acquisitions likely affect subsequent MRI processing and comparability of its derived outcomes.

Much previous work has attempted to address the complex issue of image standardization, reviewed in (Shah et al., 2011). One of the most common approaches to image harmonization is *histogram matching*, in which the intensity profile of the new image is altered to resemble that of an image template constructed from training subjects (Nyul et al., 2000). However, the histogram matching does not entirely cancel the effects of the intensity mismatches (Roy et al., 2013) and often fails to preserve biological characteristics (Fortin et al., 2016; Wrobel et al., 2020). Another popular approach is *contrast synthesis*, which instead of directly altering intensity values, uses a database of training images to generate a new image with a desired intensity profile (Iglesias et al., 2013; Roy et al., 2013). The significant drawbacks of this approach are computational demands and the need for an extensive training dataset. It has been shown to be beneficial for some intermediate processing steps such as registration but not for segmenting brain structures (Iglesias et al., 2013).

A novel intensity unit normalization method, called White Stripe (Shinohara et al., 2014), has been developed to bring raw image intensities to a biologically interpretable scale. The method applies a z-score transformation on image intensities based on the parameters estimated from the normal-appearing white matter (NAWM). The use of NAWN makes the method useful for many studies of brain abnormalities, such as multiple sclerosis lesions. While the method makes the white matter (WM) comparable across subjects, the residual across-subject intensity variations are still present in the cerebrospinal fluid (CSF) and grey matter (GM). Therefore, the method is only helpful for the intensity unit normalization.

To correct between-scan effects, the *Removal of Artificial Voxel Effect by Linear regression* (*RAVEL*) has been proposed (Fortin et al., 2016). The method is an extension of the White Stripe and uses White Stripe normalized images to correct for residual unwanted intensity variations across images. It has been shown to improve the replicability of the biological findings.

Recently, a multisite image harmonization by cumulative distribution function (CDF) alignment (*mica*) has been proposed (Wrobel et al., 2020). The method leverages multiple scans of the same subject to derive an intensity transformation that removes between-scan effects. Although it has been shown to reduce volumetric differences in WM and GM across sites, it is unclear how well the approach generalizes to variations in image acquisition and the impact on other brain structures.

Here we propose a Robust Intensity Distribution Alignment (RIDA) method to harmonize image intensities across visits and improve volumetric outcomes of brain structures in MR images acquired with different acquisition parameters. The present study builds on the previous work of *mica* (Wrobel et al., 2020) but differs in the way the intensity transformation (mapping) is derived and used. We analyze three intra-scanner and three inter-scanner protocol variations before and after between-scan effects normalization among the same 20 participants that have been scanned on the same day. Using the term *between-scan effects*, we encapsulate residual intensity variability post-normalization (after White Stripe intensity unit normalization). This allows us to systematically measure segmentation differences due to variations in acquisition parameters within and between scanners and to what degree these can be reduced by applying the image harmonization methods. We use RAVEL, *mica*, and RIDA to reduce between-scan effects, and volumetric outputs of the SAMSEG whole-brain segmentation (Puonti et al., 2016) to inspect the impact of different harmonization methods.

The unique dataset used in this work allows us to inspect the performance of the image harmonization under distinct circumstances: 1) the same MRI scanner but different image acquisition protocols; 2) the same imaging protocol but different MRI scanners; 3) different MRI scanners and imaging protocols. Therefore, the main objectives of this work are: 1) establish a framework for a robust image harmonization using CDFs (RIDA); 2) implement two versions of RIDA method – whole-brain and subcortical; 3) evaluate the performance of image harmonization methods within- and between-scanners. We also show how between-scan normalization methods work with different bias field correction methods, namely N3, N4, and the one estimated as part of the SAMSEG’s processing stream.

## 2. Material and methods

### 2.1. Datasets

A total of 20 cognitively healthy participants (18 females, age range from 20 to 36 years, mean =

27.3 years, sd = 4.5 years) were scanned using three models of Siemens MRI scanners (Siemens Medical Solutions, Erlangen, Germany): 1.5T Avanto, 3T Skyra, and 3T Prisma, at Rikshospitalet, Oslo University Hospital. The project was approved by the Regional Ethical Committee, region South, and all participants gave informed consent. For each participant, two different T1w scans were acquired from each scanner on the same day. Table 1 summarizes MRI T1w pulse sequence parameters for each scanner and outlines the comparisons of interest used in this work. For the optimal comparability between two acquisitions within each scanner, the participants were not repositioned, and one T1w acquisition immediately followed the other.

**Table 1.**
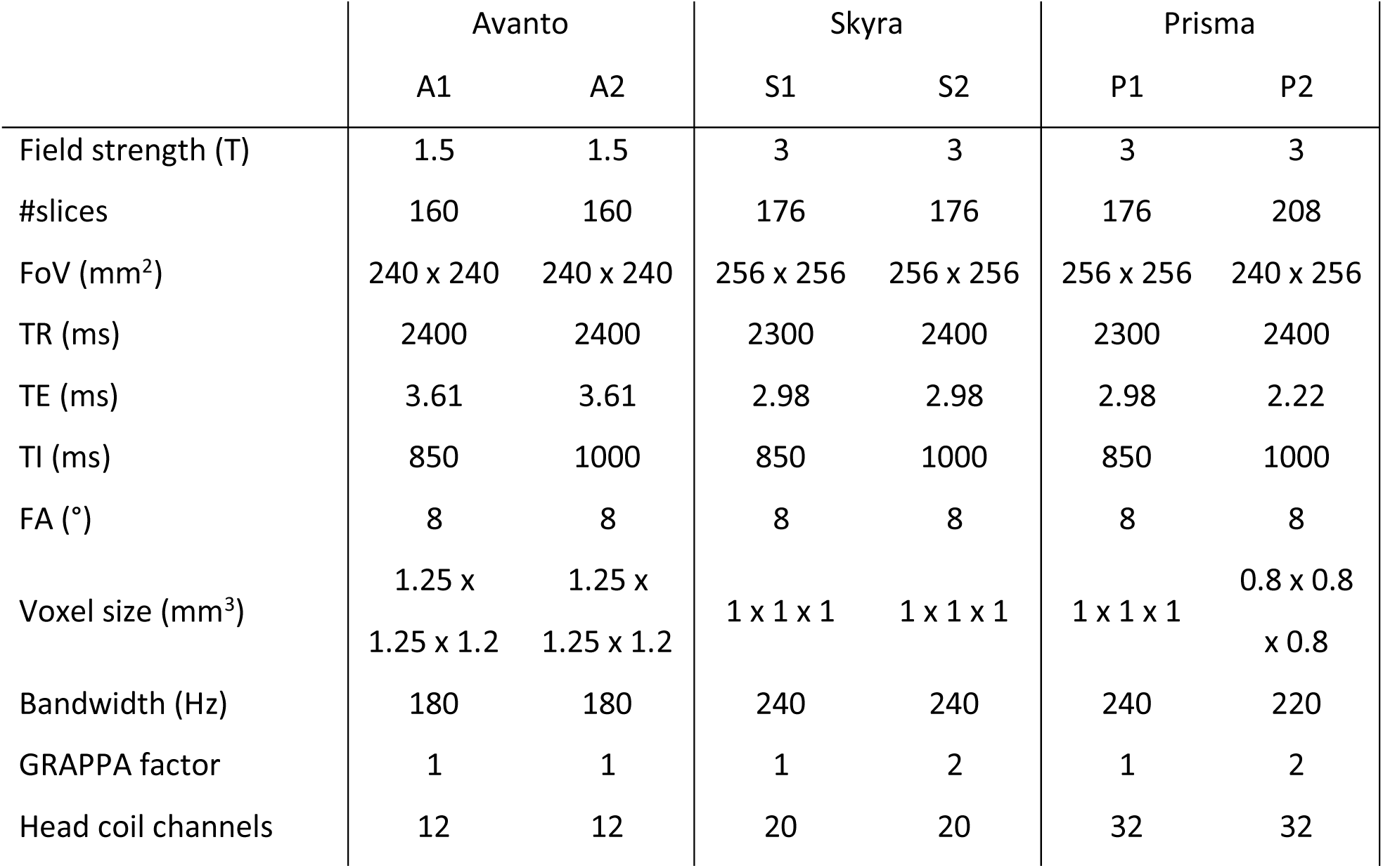
A summary of MRI T1w acquisition parameters used for the Siemens Avanto, Skyra, and Prisma scanners. The between-scan effects normalization methods are applied to all three intrascanner comparisons (A1 vs. A2, S1 vs. S2, P1 vs. P2) and a subset of inter-scanner comparisons (A2 vs. S1, A2 vs. P2, S1 vs. P1). The selected inter-scanner comparisons are particularly interesting as most of the longitudinal data at our research center has been acquired with the selected scanner changes between visits.

### 2.2. MRI pre-processing

First, due to the non-linearity of the magnetic fields from the imaging gradient coils, we preprocessed the images to reduce the geometrical variability of the same participant’s brains between different MRI scanners. We obtained scanner-specific spherical harmonics expansions representing the gradient coils (Jovicich et al., 2006).

Second, we resampled the images to the so-called *conformed* space (256×256×256 image dimensions and 1×1×1 mm^3^ isotropic voxels) using cubic interpolation and corrected for the intensity inhomogeneities (also known as a bias field) that are caused by the coil sensitivity variations as well as regional variations in brain tissue magnetic susceptibility. The bias field might not only hamper the segmentation of brain structures but also is a significant source of between-scan effects, especially when dealing with images from two different scanners. To address the sensitivity of the bias field correction to the image harmonization, we considered three bias field correction methods: N3 (Sled et al., 1998), N4 (Tustison et al., 2010), and SAMSEG’s (Puonti et al., 2016). The bias field in SAMSEG is modeled as a linear combination of spatially smooth basis functions (Van Leemput et al., 1999) and is estimated simultaneously with the brain segmentation. The approach is closely related to the unified segmentation framework described in (Ashburner and Friston, 2005).

Finally, we normalized each image using the White Stripe method (Shinohara et al., 2014). The normalization is based on the normal-appearing white matter (NAWM) and scales image intensities so that white matter becomes comparable across images. Instead of z-transforming the images initially, we scaled the intensities so that the mean NAWM intensity value was 110. We refer to these images as WM-normalized as the method aligns WM intensity distributions between images. However, residual across-subject variability is still present in the CSF and grey matter.

### 2.3. Between-scan effects normalization

Here we describe three approaches to address the issue of between-scan effects: RAVEL, *mica*, and RIDA. RIDA and *mica* are similar and based on the same underlying principle – the alignment of two cumulative distributions of voxel intensities. Both methods are expected to work best when two different images of the same subject are acquired in a short time interval. The differences between images would be attributed to the between-scan effects rather than biological differences. On the other hand, RAVEL utilizes all study images to estimate unwanted variability from a control region and perform voxel-wise intensity corrections using a linear regression model. The main advantage of RAVEL is that it does not require images of the same subject to be acquired in a short time interval and can also be of different subjects.

#### 2.3.1. RAVEL

RAVEL is a tool for removing scan effects that utilize all images in the study to leverage information about unwanted variability after intensity normalization (White Stripe). Each image of the study is non-linearly registered to a standard template to allow voxel-wise linear models and estimate the unwanted variation component from regions of the brain that are not expected to be associated with the clinical covariate of interest.

Here, we used an R package https://github.com/Jfortin1/RAVEL to perform RAVEL and preprocessed images described in Section 2.2 as inputs to RAVEL. Briefly, the unwanted variation component was estimated from a control region where voxels were labeled as a cerebrospinal fluid (CSF) for all images. These intensities are unassociated with disease status and other clinical covariates (Luoma et al., 1993). To get the corrected intensity values back into the native space, we applied the back-transformation from the template to the native space. To be consistent with the other two normalization methods, we did not use any additional covariates, which is possible by RAVEL.

#### 2.3.2. mica

Multisite image harmonization by CDF alignment (*mica*) harmonizes images by aligning cumulative distribution functions (CDFs) of the voxel intensities (Wrobel et al., 2020). Here, it estimates a nonlinear, monotonically increasing transformation of the voxel intensity values in one scan (“source”) such that the resulting intensity CDF perfectly matches the intensity CDF from a second (“target”) scan. These intensity transformations called warping functions, define a one-to-one mapping between intensity values from a source scan to corresponding intensity values from the target scan. The CDF alignment occurs at the subject level, meaning that two images of the same subject collected in close succession are needed. The warping function is estimated for the whole brain and does not require spatial normalization (registration).

Here, we used an R package https://github.com/julia-wrobel/mica to perform *mica* image harmonization. Before running mica, we performed ROBEX skull-stripping (Iglesias et al., 2011) on the pre-processed images described in Section 2.2. Then, we computed CDFs independently for each image and derived warping functions from CDF alignment to generate harmonized images.

#### 2.3.3. RIDA

The goal of the image harmonization framework based on CDFs is to transform the voxel intensities of one image to match the intensities of another image. Therefore, throughout the descriptions below, we assume that we operate on two MRI T1w images of the same participant – a source and a target – that have been acquired with distinct acquisition parameters, either within the same or across different scanners. Any technical variability resulting from different protocol implementations or differences between scanners are referred to as between-scan effects.

The underlying principle of the Robust Intensity Distribution Alignment (RIDA) is the same as for *mica* – align two intensity distributions and derive a warping function. However, several important improvements and considerations make RIDA more robust and reliable for image harmonization. Here, we present a framework with two parts: an additional image pre-processing required to harness the potential of close succession scans and a robust intensity distribution alignment approach.

##### Pre-processing

We used pre-processed images described in Section 2.2 to further pre-process for the RIDA. First, we co-registered two images to their halfway space using FreeSurfer’s *mri_robust_register* program (Reuter et al., 2010). The program computes an inverse consistent registration of two volumes via an iterative symmetric alignment of the two images. It uses a method based on robust statistics to detect and remove outliers from the registration. Second, we non-linearly registered the source image to the target image using ANTs diffeomorphic registration (Avants et al., 2011). This allowed for better spatial normalization and voxel-wise intensity comparisons. Third, we performed whole-brain segmentation of both images using SAMSEG to label brains into different regions later used for the intensity mappings. Finally, we White Stripe normalized both images based on the common NAWM mask, which is the intersection of two NAWM masks – one for each image. We then used the pre-processed T1w images and their corresponding whole-brain segmentation for the robust intensity distribution alignment described below.

##### A robust intensity distribution alignment

Given a pair of pre-processed images of the same subject - a source (*I*_*s*_) and a target (*I*_*t*_) - we computed two CDFs, namely *cdf*_*s*_ and *cdf*_*t*_, within a region of interest (*ROI*) which was selected from the whole brain segmentation. Next, we derived intensity mappings from *cdf*_*s*_ to *cdf*_*t*_, and from *cdf*_*t*_ to *cdf*_*s*_ at pre-defined intensity query points *x* (from 0 to 150, n = 300) by using 1D linear interpolation. Each such mapping defined a non-linear transformation (a lookup table) where a particular intensity value of one image was mapped to the intensity value of another image. In other words, we aligned two intensity distributions both ways.

Finally, to make this transformation unbiased towards the selected target image, we found an inverse consistent transformation by calculating the inverse of *cdf*_*t*_ to *cdf*_*s*_ mapping and averaging it with the *cdf*_*s*_ to *cdf*_*t*_ mapping, see Fig. 1. The derived transformation then was used to transform the intensities of the source image *I*_*s*_ (which was still non-linearly registered to the target image), so it became more like the target image, see Fig. 2.

**Fig. 1.**
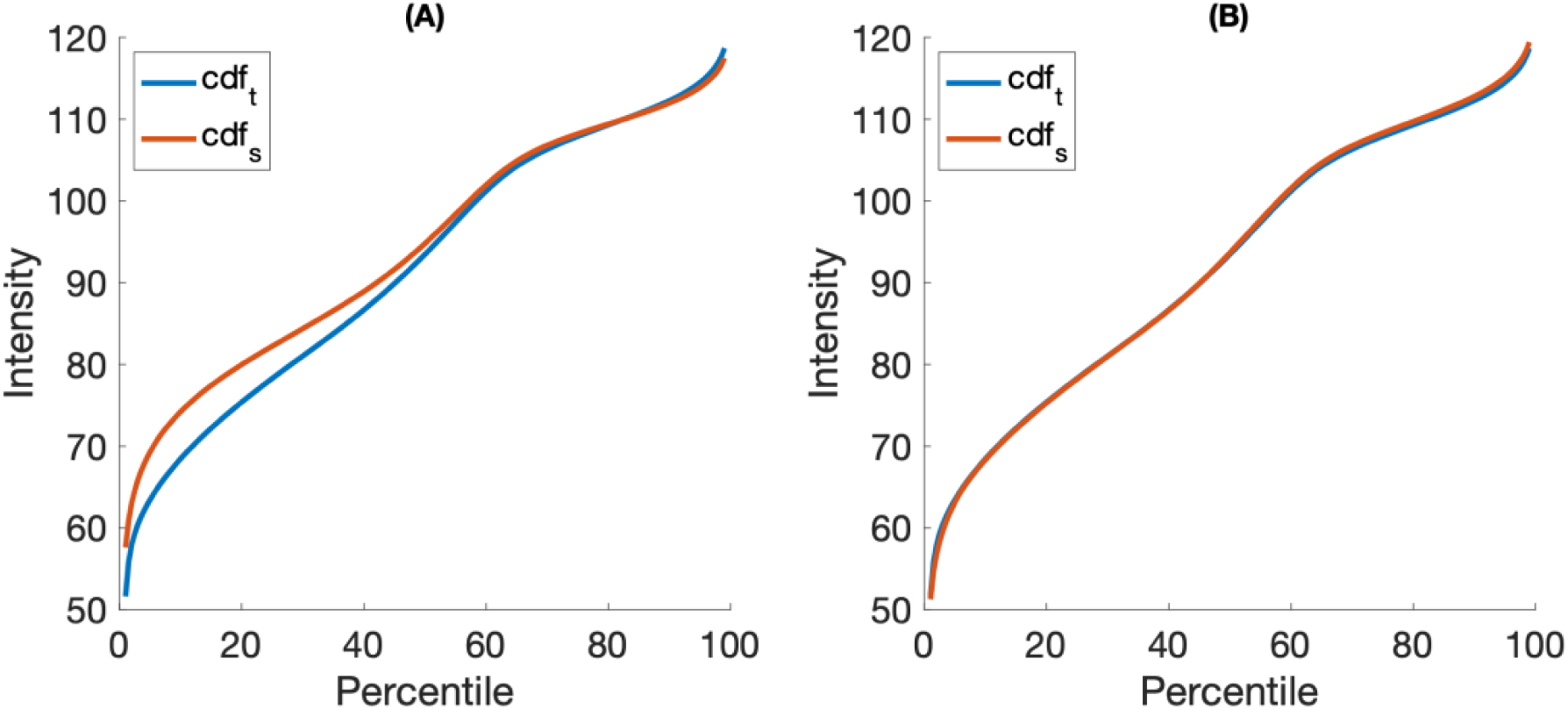
An example of two CDFs and their alignment. Figure (A) shows the intensity profiles derived from two different T1w images of the same participant – a source (cdf_s_) and a target (cdf_t_). Figure (B) shows the intensity profiles after the intensity normalization.

**Fig. 2.**
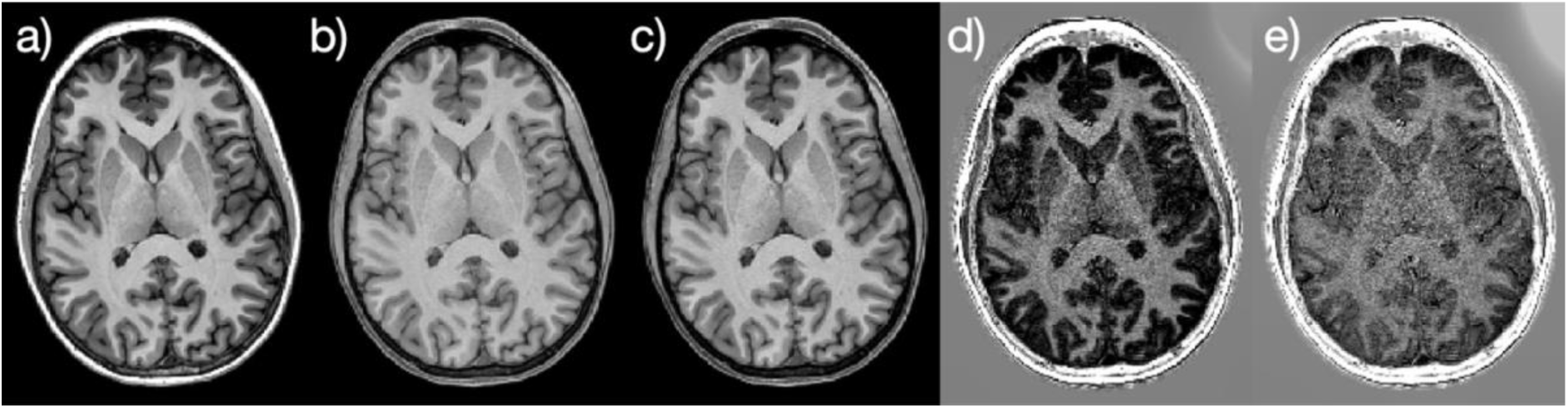
An example of intensity harmonization by CDF alignment. The images represent a) a target image; b) a source image; a source image normalized to match the intensities of the target image; d) the original voxel-wise differences between target and source images; e) voxel-wise differences between target and normalized source images. Images d) and e) are of the same intensity scale.

We repeated the same procedure twice, but we excluded potential outliers within the ROI before calculating CDFs. These outliers were characterized as voxels with much larger voxel-wise differences in intensities compared to the rest of the voxel-wise differences within the *ROI* after intensity normalization of the source scan. This can happen due to noise, poor registration, or the protocol implementations where one image can contain structures that are not present in another image, for example, blood vessels. Therefore, to reduce the effects of outliers, we excluded voxels for which the absolute difference between images was three times larger than the median of all absolute differences within the *ROI*:

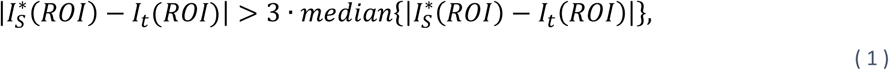

where 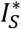 is the source scan after image harmonization. The whole procedure can be summarized as follows:

1. Calculate *cdf*_*s*_ and *cdf*_*t*_ within the selected *ROI*.
2. Derive intensity mappings from *cdf*_*s*_ to *cdf*_*t*_, and from *cdf*_*t*_ to *cdf*_*s*_.
3. Calculate the inverse of *cdf*_*t*_ to *cdf*_*s*_ mapping and average it with the *cdf*_*s*_ to *cdf*_*t*_ mapping.
4. Apply the average intensity mapping to the source image.
5. Update *ROI* mask by excluding voxels that match (1) criterion.
6. Repeat steps 1-5 two more times.

We used the final derived transformation (after three iterations) to reduce between-scan effects by transforming the intensities of the source image in its *native space* (before spatial normalization). Still, we derived the intensity mapping from two images spatially normalized by the non-linear registration to allow the robust exclusion of outlier voxels, see Fig. 3.

**Fig. 3.**
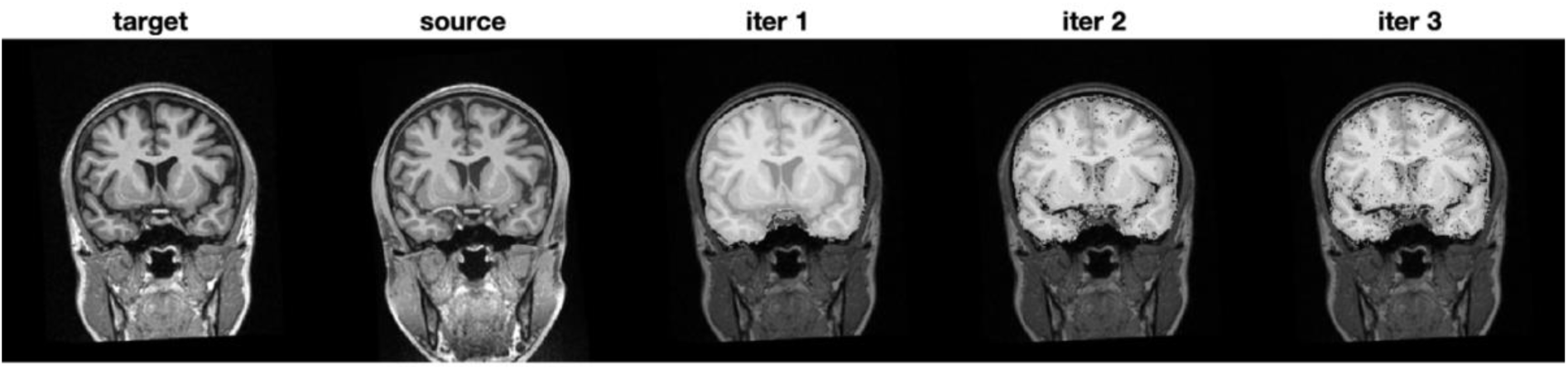
Removal of outlier voxels. The source and target images are spatially normalized by non-linear registration to improve voxel-wise correspondence. White overlayed masks (iter 1, iter 2, and iter 3) represent brain voxels from which the intensity mapping is derived at each iteration. The target image’s initial brain mask (iter 1) gets refined by excluding outlier voxels.

The application of the framework largely depends on the selected ROI. Consequently, we present two ROI-specific applications of the framework, namely whole-brain and subcortical intensity harmonization.

##### Whole-brain intensity harmonization

The whole-brain image harmonization aims to transform intensity values of all brain regions at once. Therefore, a natural choice of ROI for the framework is a brain mask that covers the entire brain area. To be more specific, given two pre-processed MRI images and their whole-brain segmentation (as described in the MRI pre-processing section), we found whole-brain masks for each image which we used to derive a robust intensity mapping. Then, we applied the intensity transformation to the pre-processed source image in the native space (before spatial normalization). This resulted in a corrected image where voxel intensities were altered to match the intensities of the target image.

##### Subcortical intensity harmonization

We also defined an ROI that only covered the subcortical brain structures and did not include cortical regions. This ROI directly corresponded to the brain structures of interest analyzed in this paper. This ROI selection was used to test whether cortical intensity values (and their surroundings) significantly impacted the scan effects of the subcortical structures.

### 2.4. Brain segmentation

We used a fully automated whole-brain segmentation method SAMSEG (part of FreeSurfer v7.2) to label eight brain structures of interest: amygdala, caudate, hippocampus, lateral ventricles, inferior lateral ventricles, pallidum, putamen, and thalamus. Briefly, SAMSEG utilizes a mesh-based atlas and a Bayesian modeling framework to obtain volumetric segmentation without the need for skullstripping as it includes segments for the extra cerebrospinal fluid, skull, and soft tissue outside of the skull. To extract reliable segmentation results, we processed pairs of images with a longitudinal stream in SAMSEG. The longitudinal SAMSEG is based on a generative model of longitudinal data (Cerri et al., 2020; Iglesias et al., 2016). In the forward model, a subject-specific atlas is obtained by generating a random warp from the usual population atlas. Subsequently, each time point is again randomly warped from this subject-specific atlas. Bayesian inference is used to obtain the most likely segmentation, with the intermediate subject-specific atlas playing the role of latent variable in the model, whose function is to ensure that various time points have atlas warps that are similar between themselves without having to define a priori what these warps should be like.

### 2.5. Statistics

To evaluate the consistency of post-processing outcomes after image harmonization, we used the absolute symmetrized percent difference (ASPD) metric between a pair of images of each subject and the brain structure of interest:

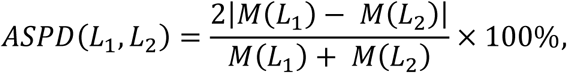

where *L*_1_ and *L*_2_ are the segmented labels of the same structure but of different images and *M(L*) is the measurement of interest calculated for that label. ASPD value closer to 0 indicates a better quantification consistency between a pair of images. We used ASPDs to evaluate differences in image-derived measurements: mean intensity, volume, and spatial overlap (Dice scores) (Dice, 1945). We separately calculated ASPD and Dice scores for the left and right hemispheric structures and then averaged.

We also conducted a three-way repeated-measures ANOVA to examine the effects of brain structure, study, and image harmonization method on volume ASPD values in 20 subjects. In addition, we used paired *t-tests* with Bonferroni multiple comparisons correction (Bland and Altman, 1995) to test whether there were significant pairwise differences among intensity harmonization approaches within each study.

All statistical analyses described above were done using R statistical software package v3.6.3 (R Core Team, 2020) and its related packages: *ggplot2* (Wickham, 2016), *ggpubr* (Kassambara, 2020), and *dplyr* (Wickham et al., 2020).

## 3. Results

### 3.1. The effect of bias field correction

Fig. 4 presents volume ASPDs for a single study, *A vs. S*, and three bias field corrections: N3, N4, and SAMSEG. The choice of the bias field correction had almost no impact (only slightly worse for N4) for RAVEL method. This was expected as RAVEL does linear regression on a voxel-wise basis. Contrary to RAVEL, *mica* and RIDA had significantly different outcomes due to the choice of the correction as both methods operate globally and include residual bias fields. This becomes evident in Fig. 5, where White Stripe normalized images, and their corresponding NAWM labels are shown. The residual bias fields after N3 and N4 hampered NAWN detection in the frontal lobe area, whereas this was not the case for SAMSEG’s bias-corrected images. Hence, for most brain structures, the intensity mappings derived from SAMSEG’s bias field corrected images showed significant improvement in the consistency of volume quantification between scanning protocols. Therefore, it was used for further analysis and results.

**Fig. 4.**
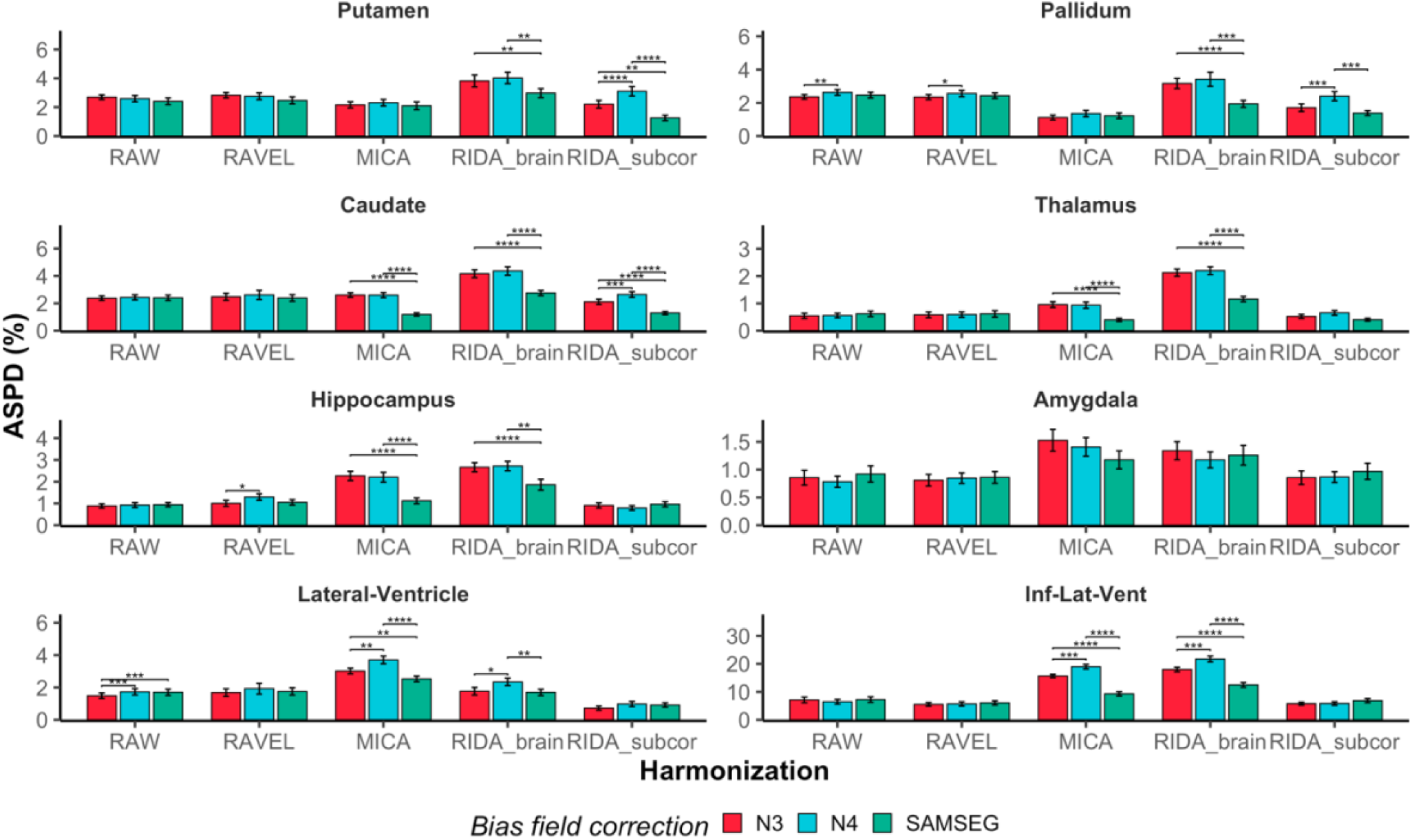
The effect of the bias field correction on volume ASPDs between the Avanto (A) and the Skyra (S) scanners. RAW indicates White Stripe normalized images used as input to the image harmonization methods: RAVEL, MICA, and RIDA (whole brain and subcortical). The vertical bars show standard errors, and significant pairwise differences are indicated by the horizontal bars with the following significance codes of the p-values: 0.0001 ‘****’, 0.001 ‘***’, 0.01 ‘**’, 0.05 ‘*’.

**Fig. 5.**
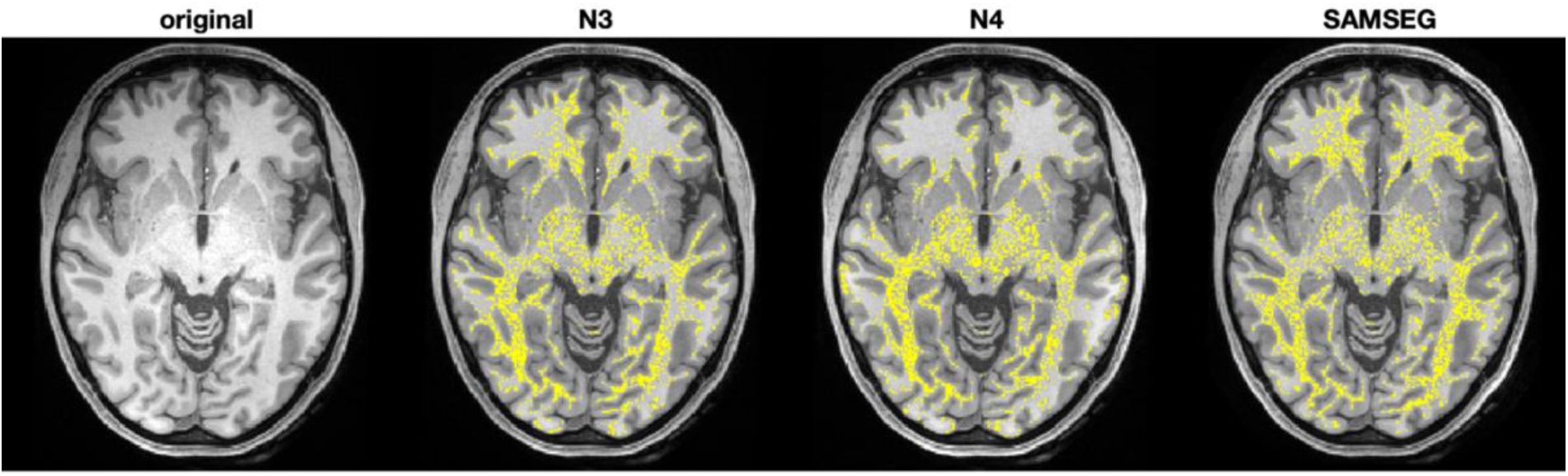
White Stripe Normal Appearing White Matter (NAWM) marked in yellow after removal of bias field. Regional intensity variations are still present in the frontal lobe area after N3 and N4 but less apparent after SAMSEG’s bias field correction, leading to a more uniform NAWM labeling.

### 3.2. *mica* vs. RIDA intensity mappings

Fig. 6 presents individual intensity mappings of 20 subjects derived from a single study, A vs. S, using *mica* and RIDA scan-effect normalization methods. More considerable inter-subject variability observed at the higher intensities is less critical since these usually correspond to non-brain regions such as the skull or blood vessels. The increased inter-subject variability was observed at the lower intensities. This is because the source and target images have an unequal number of brain voxels from which the CDFs are estimated and do not have a perfect spatial voxel-wise correspondence. The number of brain voxels varies from image to image even for the same subject due to the skull stripping, poorer spatial normalization around the cortex, and CSF which represent most of the lower intensities.

**Fig. 6.**
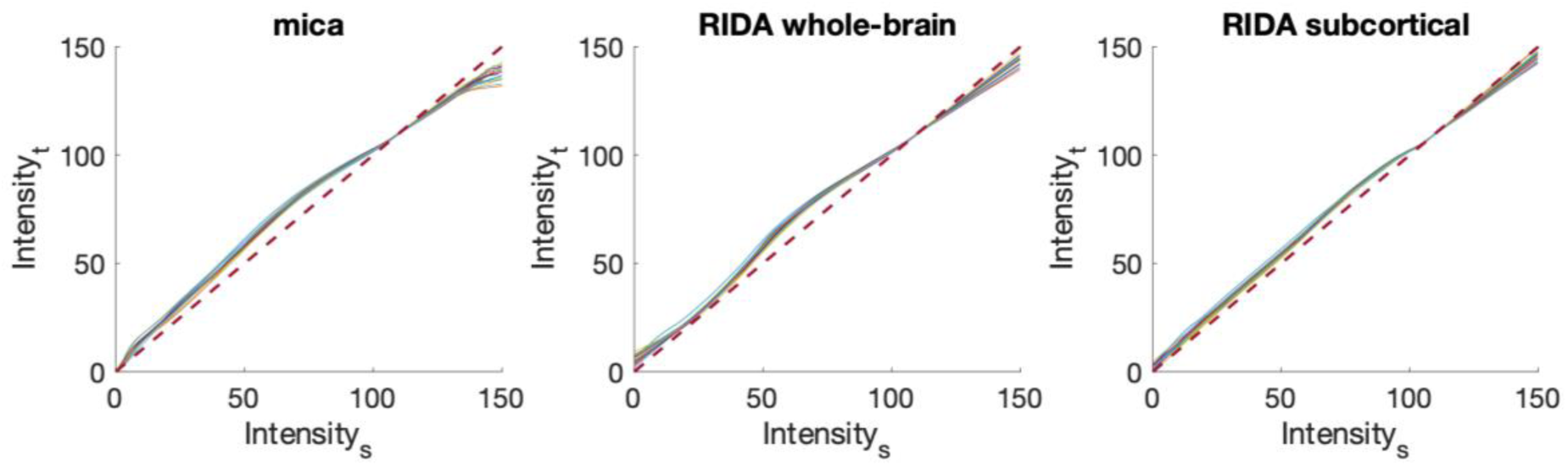
Intensity mappings of 20 subjects from a single study Avanto (A) vs. Skyra (S). Source intensities (Intensity_s_) are mapped to the target intensities (Intensity_t_). A dashed line indicates identity transformation, which has no effect between images when applied.

### 3.3. Consistency in image-derived measurements

Fig. 7 presents the mean intensity ASPDs per brain region and study. CDF-based intensity harmonization significantly reduced intensity differences between images, whereas slight improvement was observed for RAVEL. In most cases, RIDA subcortical approach provided superior results.

**Fig. 7.**
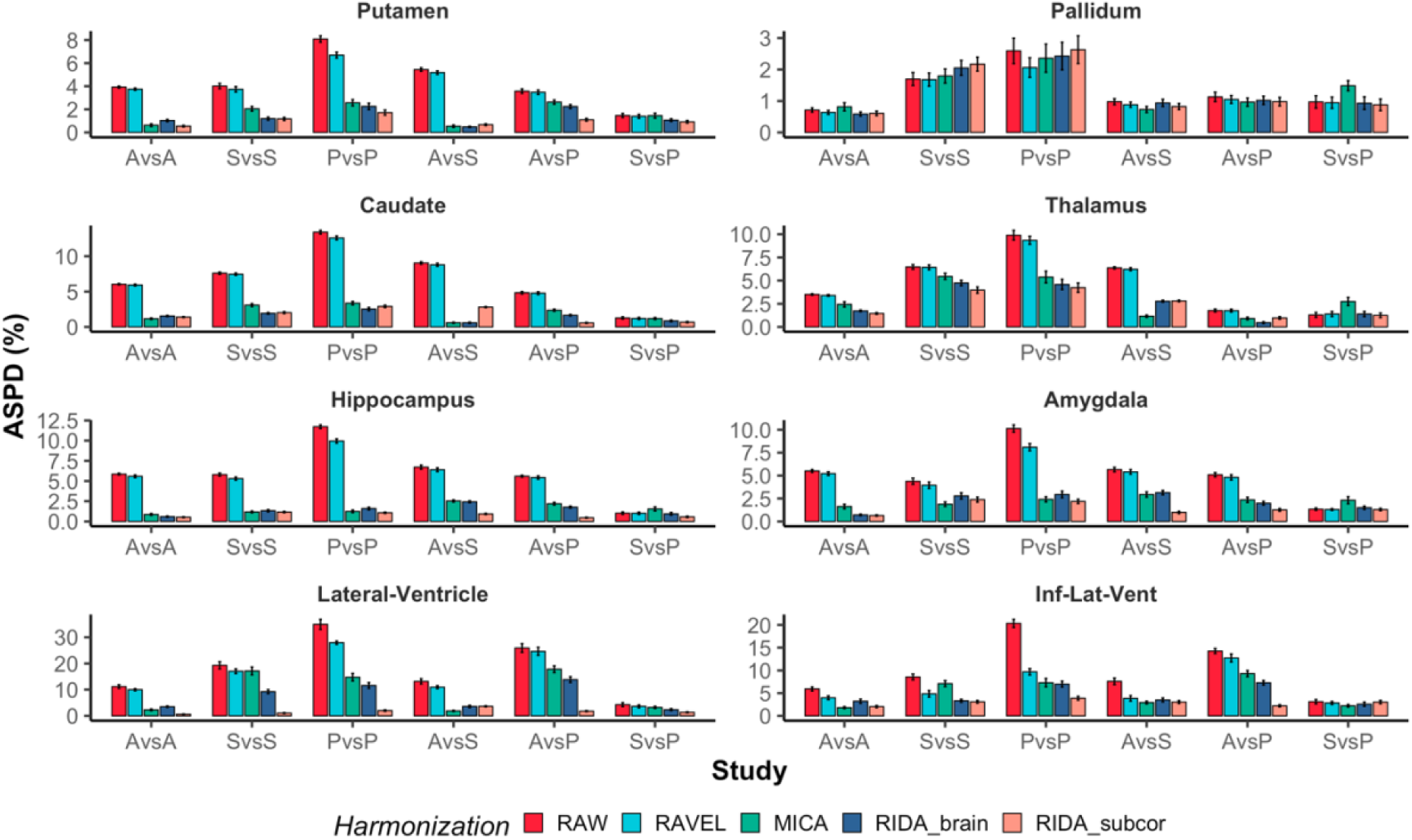
Mean intensity ASPDs per brain region. RAW indicates White Stripe normalized images used as input to the scaneffect normalization methods: RAVEL, MICA, and RIDA (whole brain and subcortical). Standard errors are shown by the vertical bars.

Fig. 8 presents volume ASPDs per brain region and study. A three-way repeated-measures ANOVA of volume ASPDs indicated a significant interaction between brain structure, study (protocol comparison), and image harmonization approach (*F*(140,4541) = 8.44, *p* < 0.0001) as well as all two-way and one-way interactions (p-values < 0.0001). Therefore, the results of the image harmonization depend on a particular setup of the pulse sequences, a structure of interest, and a normalization method.

**Fig. 8.**
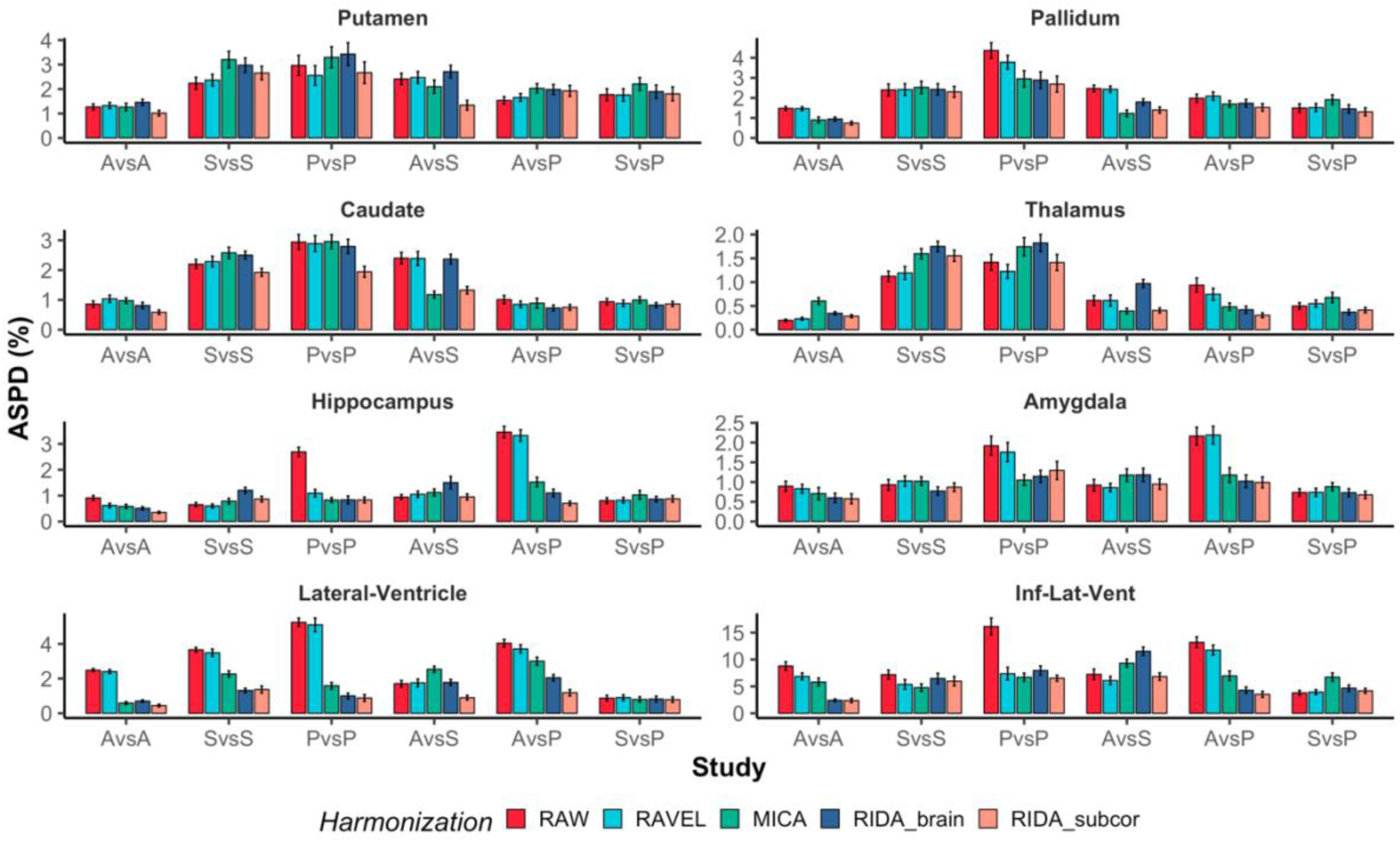
Volume ASPDs per brain structure. RAW indicates White Stripe normalized images used as input to the scan-effect normalization methods: RAVEL, MICA, and RIDA (whole brain and subcortical). Standard errors are shown by the vertical bars.

An evident advantage of the CDF-based intensity harmonization (*mica* and RIDA) was seen for ventricles, hippocampus, amygdala, and caudate. The largest improvements were observed for Prisma vs. Prisma (PvsP) and Avanto vs. Prisma (AvsP) studies. On the other hand, RAVEL showed a slight improvement compared to other approaches but did not tend to make the results worse than before the image normalization – it either stayed around the same level or slightly improved.

The performance of *mica* was comparable to the RIDA whole-brain approach, but in more cases, the latter yielded even smaller volumetric differences between the images, especially for the ventricles. RIDA subcortical-based corrections reduced initial volumetric differences more than the RIDA wholebrain corrections, for example, putamen and caudate (AvsA and AvsS), hippocampus (AvsA and AvsP), lateral ventricles (AvsA, AvsS, and AvsP). There also were cases where RIDA whole-brain corrections increased initial (uncorrected) volumetric differences. Still, subcortical-based corrections reduced it, for example, putamen (AvsA, PvsS, and AvsS), thalamus (PvsP and AvsS), and hippocampus (AvsS). Overall, the subcortical-based RIDA was superior to the whole-brain RIDA.

Numerical results of Fig. 8 are presented in the appendix (see Fig. A.1) along with the spatial overlap (Dice scores) results (see Fig. A.2). CDF-based normalization, in general, tended to increase the spatial overlap, which was already large. There was little difference in the spatial overlap between RIDA whole-brain and subcortical approaches, whereas RAVEL did not indicate any improvements.

## 4. Discussion

CDF-based image harmonization significantly improved the consistency of image-derived measurements between scanning protocols. This was particularly true for the RIDA subcortical-based transformations, which often worked better than the whole-brain corrections and did not make differences larger even in the few cases where the whole-brain corrections did. The benefit of the subcortical-based transformation comes from excluding the cortical intensity values that overlap with a range of the subcortical regions’ intensity values. This indicates that a more specific intensity transformation can provide more consistent segmentation outcomes. However, the extent to which the intensity harmonization improved the downstream results depended on several factors, including scanner/sequence comparisons and regions in the brain. Hence, while CDF-based intensity harmonization shows substantial potential to improve the consistency of image-derived measurements, the exact choices recommended for a particular study will depend on these different factors.

We described an MRI T1w image pre-processing for each image regardless of the image harmonization method applied. Notice that, in practice, any other method could be used for a particular pre-processing step, such as the bias field correction and intensity unit normalization, not necessarily those presented in this paper. We also compared the impact of the different bias field corrections and found that the bias field corrected image resulting from the SAMSEG processing yielded the most reliable results.

RAVEL is entirely data-driven and does not necessarily need multiple scans for the same subject. However, the method does not demonstrate improved consistency in image-derived measures to a similar extent as the CDF-based approach. RAVEL assumes that the space spanned by the unwanted factors estimated from the control voxels also spans the unwanted variation space for all voxels, which in this case may be violated. Another drawback is that a large study requires huge memory resources as the whole dataset must be stored in the memory and non-linear registration to the standard template is time-consuming.

RIDA and *mica* are based on the same principle – the alignment of two cumulative distributions of voxel intensities. *mica* and RIDA need specific datasets which usually are expensive to obtain or sometimes infeasible. However, given the benefits of the CDF-based harmonization, it might be worth the extra effort. In addition, *mica* does not require spatial registration (normalization) and the whole-brain segmentation as opposed to the RIDA but is less robust to the regional structure variations and does not handle outliers.

The robustness of the RIDA method comes from two improvements: non-linear diffeomorphic registration and exclusion of outliers. Thus, the correction for geometrical distortion might not be needed for close succession scans as the non-linear diffeomorphic registration can account for regional variations. This makes the method robust to geometrical differences between the scanners. The exclusion of outliers makes the procedure less sensitive to different pulse sequence implementations. The advantage of both can be observed from the results where the whole-brain RIDA generally yielded better consistency in measurement quantification.

The most notable improvements for nearly all pulse sequence comparisons were observed for the ventricles. This is likely due to the lower intensity values that are primarily present within ventricles, and do not overlap with other intensity distributions, see Fig. 9. It also confirms the observation from Fischl et al. (2002) that the hippocampus and amygdala have almost completely overlapping intensity profiles, which necessitates using probability information in addition to intensity in segmenting these structures. Nevertheless, whole-brain and subcortical RIDA corrections significantly reduced the differences between 1.5T Avanto and 3T Prisma, and between the two Prisma sequences, in similar ways for the hippocampus and amygdala. This shows that the correction can affect the segmentation of structures with the same intensity profiles in similar ways. This further underscores the point that optimal correction procedure depends on multiple parameters, including sequence, scanner, and brain region of interest.

**Fig. 9.**
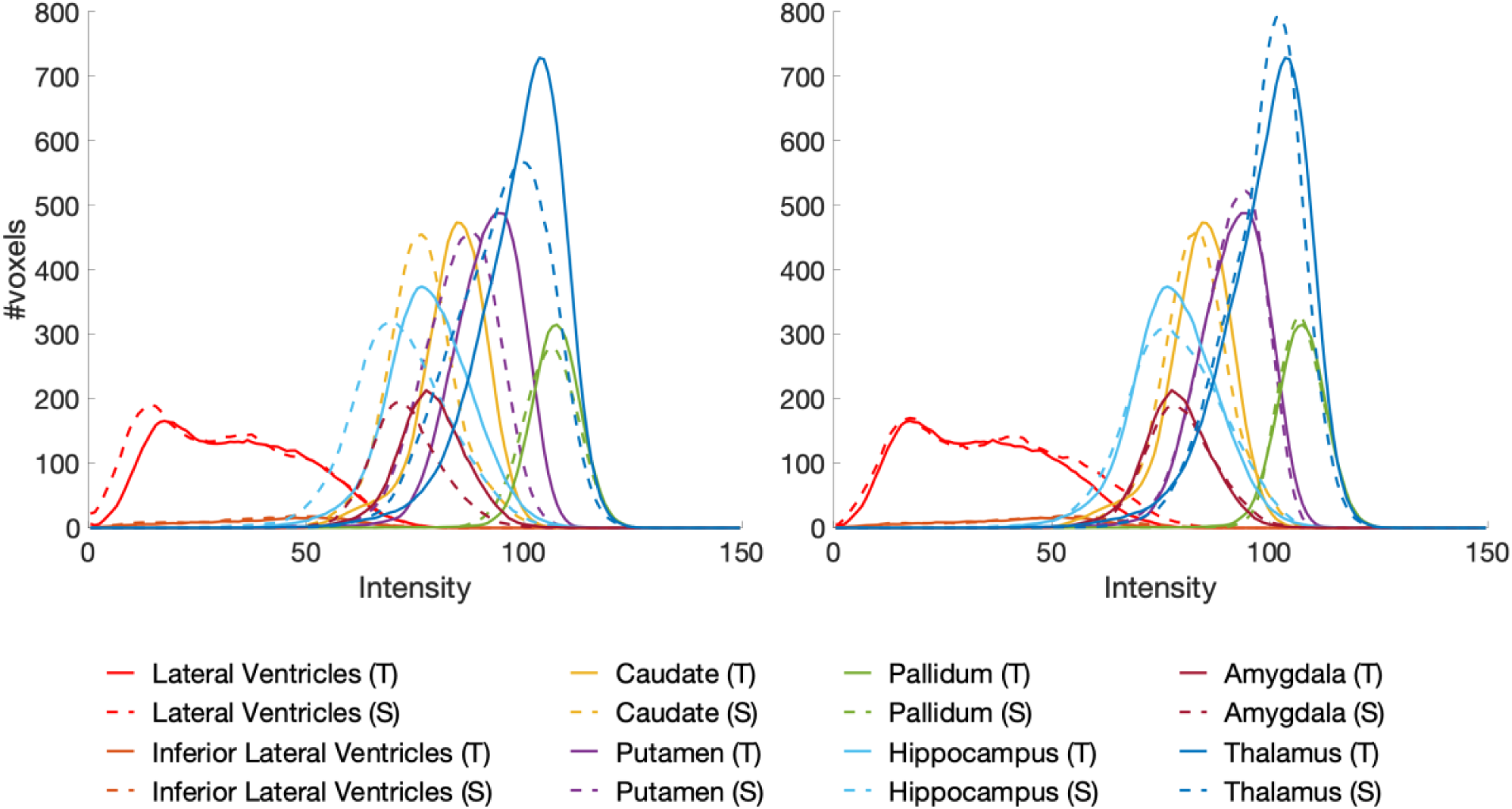
An example of the intensity distributions before (left) and after (right) RIDA intensity harmonization for 1.5T Avanto vs. 3T Skyra comparison for a single participant. The source (S) distributions are displayed as dashed lines and the target (T) distributions as solid lines. Lower intensities are primarily present within ventricles and almost do not overlap with other subcortical structures.

The outcomes for Skyra vs. Prisma (S vs. P) study, which represents the same pulse sequence but different scanners, did not show any improvements for neither of the image harmonization methods. The individual intensity mappings shown in Fig. 10 indicate that neither *mica* nor RIDA was sensitive enough to model the between-scanner differences (scanner effects) - the mappings were close to identity transformations meaning that small intensity changes were done. Interestingly, while the differences in estimated volumes tended to be relatively small for this comparison – 3T Skyra vs. 3T Prisma with the same sequence parameters – the correction procedure was often able to reduce the initially more considerable differences between the other scanners/sequences down to a level comparable with the Prisma – Skyra same sequence differences. However, while the correction procedure could not reduce this between-scanner difference in the present study, the procedure still warrants further investigation for such corrections. Our comparisons were between Siemens scanners, and it is possible that the correction would reduce differences between scanners from different vendors.

**Fig. 10.**
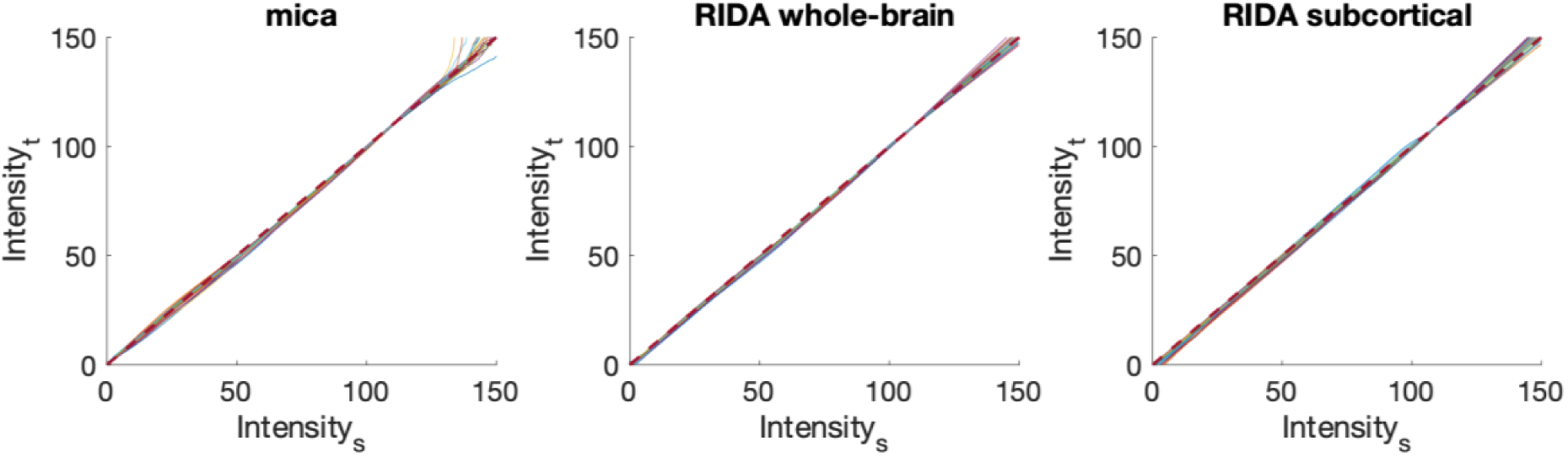
Intensity mappings of 20 subjects from a single study Skyra (S) vs. Prisma (P) represent the same MRI acquisition protocol but different scanners. Source intensities (Intensity_s_) are mapped to the target intensities (Intensity_t_). A dashed line indicates identity transformation, which has no effect between images when applied.

In the intensity harmonization by CDF (either *mica* or RIDA), we assume that we have a pair of images – a source and a target – for the same participant scanned within a short time interval. This is necessary to derive a reliable intensity mapping from one image to another. In practice, however, we would only acquire a small number of paired scans to derive a transformation between two specific MRI acquisitions, either within or between the scanners. The rest of the images would be acquired by one of the acquisitions. Therefore, to reflect a more realistic scenario, we would need to derive an average intensity mapping from a training dataset and apply it to a new independent test dataset. In this work, however, we derived and used intensity mappings under “perfect” conditions meaning that each subject had a unique intensity transformation. This yielded overly too optimistic outcomes.

Nevertheless, suppose each subject’s unique intensity mapping cannot improve consistency in between-scan measurements. In that case, it is unreasonable to expect it to work any better in a more realistic scenario. Therefore, this empirical study gives us a glimpse of the performance of each harmonization method, allowing us to compare inter-individual variations. In general, as seen from Fig. 6, the observed individual intensity mappings of *mica* and RIDA indicated low inter-individual variability meaning that a reliable average transformation could be derived.

In this work, we have only analyzed the subcortical structures. The next step would be to inspect how the CDF-based harmonization works for the surface-based analysis where, for example, the cortical thickness measures would be of interest. Moreover, instead of deriving the exact transformation for each subject, we must inspect how well the RIDA generalizes. We need to derive an average intensity mapping from a sample of subjects and apply it to other subjects. Furthermore, we have seen that using a targeted intensity transformation (subcortical-based) yielded better results than the general approach (whole-brain). Therefore, this indicates a potential for further improvements, possibly the application of convolutional neural networks, which might help avoid the overlapping intensity distributions of different structures, which in turn causes favorable outcomes for some structures but not others.

## 5. Conclusions

The intensity harmonization by cumulative distribution function can be useful for increasing the intra- and inter-scanner reproducibility of subcortical segmentation in cases of MRI pulse sequence parameter variations. However, the outcomes vary between subcortical structures and depend on a particular pulse sequence comparison. Therefore, a pilot study tailored to the sequences and scanners in question is recommended before applying the RIDA framework to a large-scale study, which can be expensive or infeasible due to the necessary condition of participants being scanned in close succession. However, in studies with a limited number of scanners and sequences, or longitudinal studies that for various reasons need to change scanners between follow-up examinations, it may be worth the effort to collect a set of multiple scans of the same participants using each of the sites/sequences that are part of the study. That will enable selecting the optimal correction method to reduce differences. Suppose no such paired scans are available and cannot be obtained. In that case, the present results still suggest that 1) performing corrections yield smaller differences than not performing corrections, and 2) the subcortical-based intensity correction (RIDA subcortical) tends to yield smaller differences than the whole-brain correction.

## Acknowledgments

The present research was funded by a grant from Helse-Sør Øst (grant number 2018009), the European Research Council under grant agreements 283634, 725025 (to A.M.F.), 313440 (to K.B.W.), as well as the Norwegian Research Council (to A.M.F., K.B.W.). Support for this research was provided in part by the BRAIN Initiative Cell Census Network grant U01MH117023, the National Institute for Biomedical Imaging and Bioengineering (P41EB015896, 1R01EB023281, R01EB006758, R21EB018907, R01EB019956, P41EB030006), the National Institute on Aging (1R56AG064027, 1R01AG064027, 5R01AG008122, R01AG016495, 1R01AG070988), the National Institute of Mental Health (R01 MH123195, R01 MH121885, 1RF1MH123195), the National Institute for Neurological Disorders and Stroke (R01NS0525851, R21NS072652, R01NS070963, R01NS083534, 5U01NS086625,5U24NS10059103, R01NS105820, R01NS112161), and was made possible by the resources provided by Shared Instrumentation Grants 1S10RR023401, 1S10RR019307, and 1S10RR023043. Additional support was provided by the NIH Blueprint for Neuroscience Research (5U01-MH093765), part of the multi-institutional Human Connectome Project. In addition, BF has a financial interest in CorticoMetrics, a company whose medical pursuits focus on brain imaging and measurement technologies. BF’s interests were reviewed and are managed by Massachusetts General Hospital and Partners HealthCare in accordance with their conflict of interest policies.

## Data and Code Availability

The raw LCBC MRI data supporting the results of the current study may be available upon reasonable request, given appropriate ethical data protection approvals and data sharing agreements. Requests for the raw MRI data can be submitted to the last author Anders M Fjell. Fully-open raw data availability is restricted as participants have not consented to share their data publicly. All data preprocessing and analysis code will be available at https://github.com/LCBC-UiO upon acceptance of the manuscript.

## Appendix A

**Fig. A.1.**
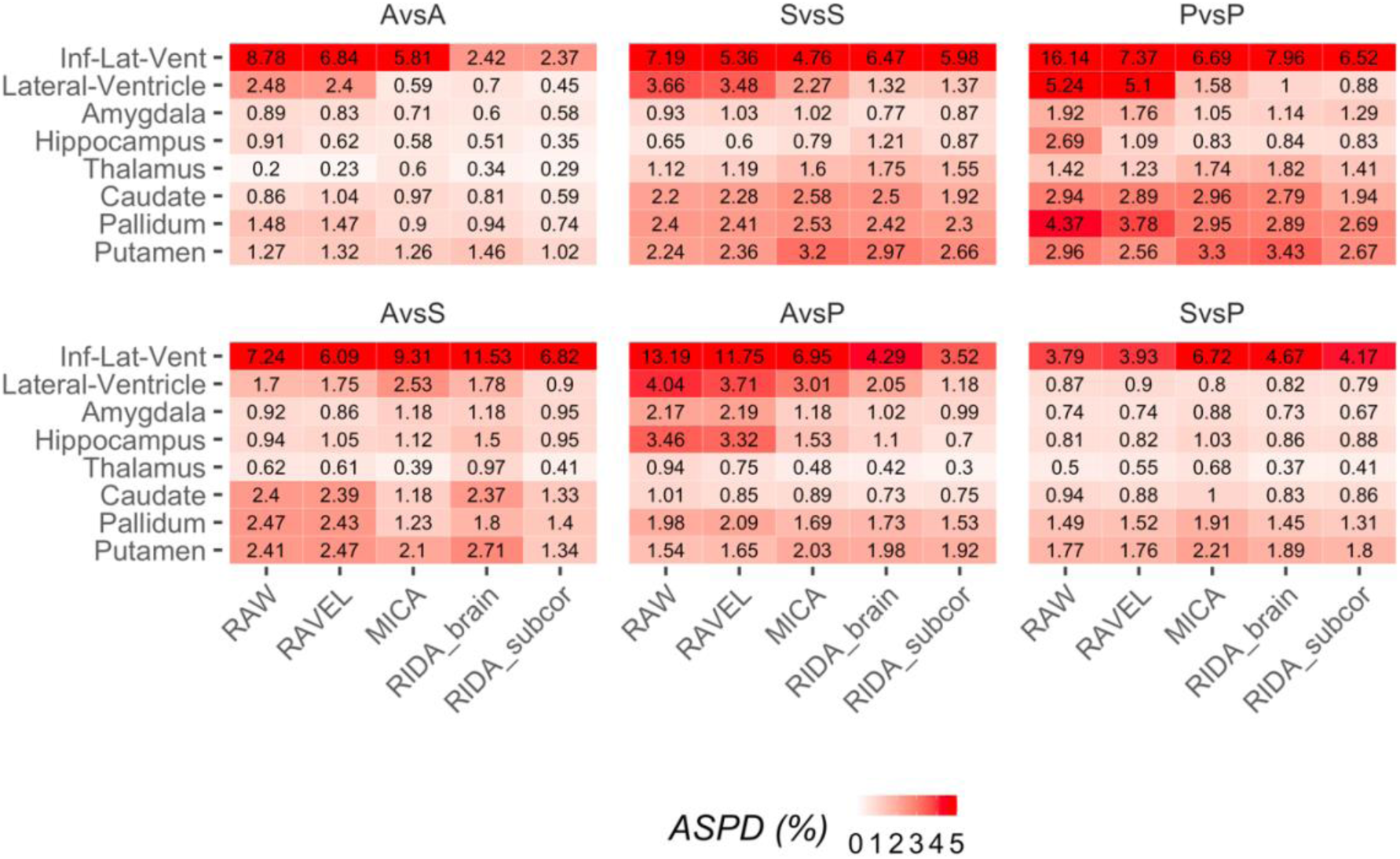
A heat map of the numerical results of the mean volume ASPDs per brain structure and study. It presents volumetric differences before (RAW) and after scan-effect harmonization (RAVEL, MICA, RIDA brain, and RIDA subcortical). To improve readability, the colors of ASPD values greater than five are ‘saturated’ and shown with the same hue.

**Fig. A.2.**
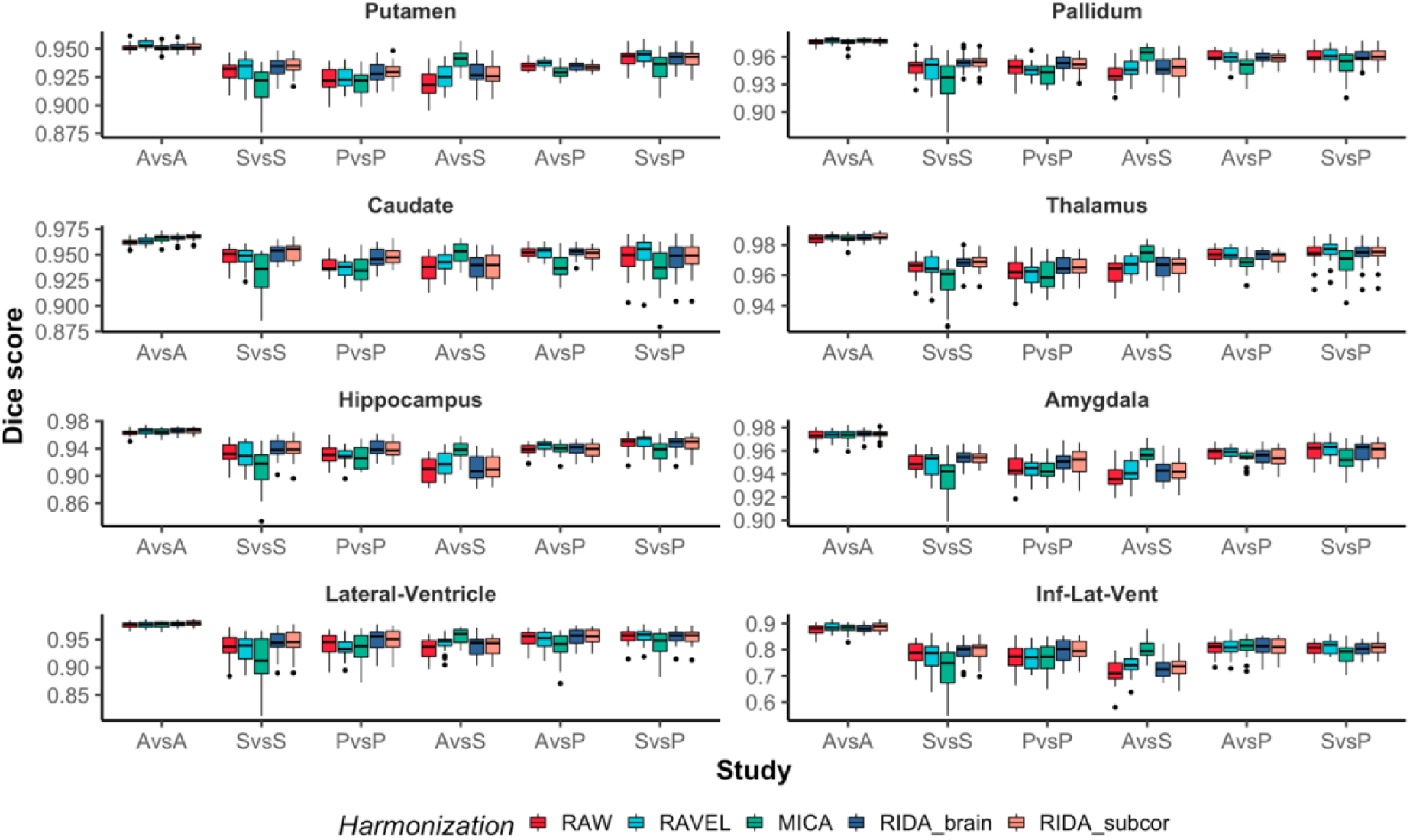
Box plots of Dice scores per brain structure and study. RAW indicates White Stripe normalized images used as input to the scan-effect normalization methods: RAVEL, MICA, and RIDA (whole brain and subcortical).

## Literature

Ashburner, J., Friston, K.J., 2005. Unified segmentation. Neuroimage 26, 839–851. https://doi.org/10.1016/j.neuroimage.2005.02.018

Avants, B.B., Tustison, N.J., Song, G., Cook, P.A., Klein, A., Gee, J.C., 2011. A reproducible evaluation of ANTs similarity metric performance in brain image registration. Neuroimage 54, 2033–2044. https://doi.org/10.1016/j.neuroimage.2010.09.025

Bland, J.M., Altman, D.G., 1995. Multiple significance tests: the Bonferroni method. BMJ 310, 170.

Cerri, S., Hoopes, A., Greve, D.N., Mühlau, M., Van Leemput, K., 2020. A Longitudinal Method for Simultaneous Whole-Brain and Lesion Segmentation in Multiple Sclerosis. 2008.05117 [cs, eess] 12449, 119–128. https://doi.org/10.1007/978-3-030-66843-3_12

Dice, L.R., 1945. Measures of the Amount of Ecologic Association Between Species. Ecology 26, 297–302. https://doi.org/10.2307/1932409

Fortin, J.-P., Sweeney, E.M., Muschelli, J., Crainiceanu, C.M., Shinohara, R.T., 2016. Removing intersubject technical variability in magnetic resonance imaging studies. Neuroimage 132, 198–212. https://doi.org/10.1016/j.neuroimage.2016.02.036

Han, X., Jovicich, J., Salat, D., van der Kouwe, A., Quinn, B., Czanner, S., Busa, E., Pacheco, J., Albert, M., Killiany, R., Maguire, P., Rosas, D., Makris, N., Dale, A., Dickerson, B., Fischl, B., 2006. Reliability of MRI-derived measurements of human cerebral cortical thickness: The effects of field strength, scanner upgrade and manufacturer. NeuroImage 32, 180–194. https://doi.org/10.1016/j.neuroimage.2006.02.051

Iglesias, J.E., Konukoglu, E., Zikic, D., Glocker, B., Van Leemput, K., Fischl, B., 2013. Is Synthesizing MRI Contrast Useful for Inter-modality Analysis?, in: Salinesi, C., Norrie, M.C., Pastor, Ó. (Eds.), Advanced Information Systems Engineering, Lecture Notes in Computer Science. Springer Berlin Heidelberg, Berlin, Heidelberg, pp. 631–638. https://doi.org/10.1007/978-3-642-40811-3_79

Iglesias, J.E., Liu, C.-Y., Thompson, P.M., Tu, Z., 2011. Robust brain extraction across datasets and comparison with publicly available methods. IEEE Trans Med Imaging 30, 1617–1634. https://doi.org/10.1109/tmi.2011.2138152

Iglesias, J.E., Van Leemput, K., Augustinack, J., Insausti, R., Fischl, B., Reuter, M., 2016. Bayesian longitudinal segmentation of hippocampal substructures in brain MRI using subject-specific atlases. NeuroImage 141, 542–555. https://doi.org/10.1016/j.neuroimage.2016.07.020

Jovicich, J., Czanner, S., Greve, D., Haley, E., van der Kouwe, A., Gollub, R., Kennedy, D., Schmitt, F., Brown, G., MacFall, J., Fischl, B., Dale, A., 2006. Reliability in multi-site structural MRI studies: Effects of gradient non-linearity correction on phantom and human data. NeuroImage 30, 436–443. https://doi.org/10.1016/j.neuroimage.2005.09.046

Jovicich, J., Czanner, S., Han, X., Salat, D., van der Kouwe, A., Quinn, B., Pacheco, J., Albert, M., Killiany, R., Blacker, D., 2009. MRI-derived measurements of human subcortical, ventricular and intracranial brain volumes: Reliability effects of scan sessions, acquisition sequences, data analyses, scanner upgrade, scanner vendors and field strengths. NeuroImage 46, 177–192. https://doi.org/10.1016/j.neuroimage.2009.02.010

Kassambara, A., 2020. ggpubr: “ggplot2” Based Publication Ready Plots.

Luoma, K., Raininko, R., Nummi, P., Luukkonen, R., 1993. Is the signal intensity of cerebrospinal fluid constant? Intensity measurements with high and low field magnetic resonance imagers. Magnetic Resonance Imaging 11, 549–555. https://doi.org/10.1016/0730-725X(93)90474-R

Nyul, L.G., Udupa, J.K., Zhang, X., 2000. New variants of a method of MRI scale standardization. IEEE Transactions on Medical Imaging 19, 143–150. https://doi.org/10.1109/42.836373

Puonti, O., Iglesias, J.E., Van Leemput, K., 2016. Fast and sequence-adaptive whole-brain segmentation using parametric Bayesian modeling. NeuroImage 143, 235–249. https://doi.org/10.1016/j.neuroimage.2016.09.011

R Core Team, 2020. R: A Language and Environment for Statistical Computing. R Foundation for Statistical Computing, Vienna, Austria.

Reuter, M., Rosas, H.D., Fischl, B., 2010. Highly accurate inverse consistent registration: A robust approach. NeuroImage 53, 1181–1196. https://doi.org/10.1016/j.neuroimage.2010.07.020

Roy, S., Carass, A., Prince, J.L., 2013. Magnetic Resonance Image Example Based Contrast Synthesis. IEEE Trans Med Imaging 32, 2348–2363. https://doi.org/10.1109/TMI.2013.2282126

Shah, M., Xiao, Y., Subbanna, N., Francis, S., Arnold, D.L., Collins, D.L., Arbel, T., 2011. Evaluating intensity normalization on MRIs of human brain with multiple sclerosis. Medical Image Analysis 15, 267–282. https://doi.org/10.1016/j.media.2010.12.003

Shinohara, R.T., Sweeney, E.M., Goldsmith, J., Shiee, N., Mateen, F.J., Calabresi, P.A., Jarso, S., Pham, D.L., Reich, D.S., Crainiceanu, C.M., 2014. Statistical normalization techniques for magnetic resonance imaging. NeuroImage: Clinical 6, 9–19. https://doi.org/10.1016/j.nicl.2014.08.008

Sled, J.G., Zijdenbos, A.P., Evans, A.C., 1998. A nonparametric method for automatic correction of intensity nonuniformity in MRI data. IEEE Trans Med Imaging 17, 87–97. https://doi.org/10.1109/42.668698

Tustison, N.J., Avants, B.B., Cook, P.A., Zheng, Y., Egan, A., Yushkevich, P.A., Gee, J.C., 2010. N4ITK: improved N3 bias correction. IEEE Trans Med Imaging 29, 1310–1320. https://doi.org/10.1109/TMI.2010.2046908

Van Leemput, K., Maes, F., Vandermeulen, D., Suetens, P., 1999. Automated model-based bias field correction of MR images of the brain. IEEE Trans Med Imaging 18, 885–896. https://doi.org/10.1109/42.811268

Wickham, H., 2016. ggplot2: Elegant Graphics for Data Analysis. Springer-Verlag New York.

Wickham, H., François, R., Henry, L., Müller, K., 2020. dplyr: A Grammar of Data Manipulation.

Wrobel, J., Martin, M.L., Bakshi, R., Calabresi, P.A., Elliot, M., Roalf, D., Gur, R.C., Gur, R.E., Henry, R.G., Nair, G., Oh, J., Papinutto, N., Pelletier, D., Reich, D.S., Rooney, W.D., Satterthwaite, T.D., Stern, W., Prabhakaran, K., Sicotte, N.L., Shinohara, R.T., Goldsmith, J., 2020. Intensity warping for multisite MRI harmonization. NeuroImage 223, 117242. https://doi.org/10.1016/j.neuroimage.2020.117242

